# NRF2 activators restrict coronaviruses by targeting a network involving ACE2, TMPRSS2, and XPO1

**DOI:** 10.1101/2025.02.24.639813

**Authors:** Fakhar H. Waqas, Francisco J. Zapatero-Belinchón, Madalina E. Carter-Timofte, Lisa Lasswitz, Demi van der Horst, Rebecca Möller, Julia Dahlmann, Ruth Olmer, Robert Geffers, Gisa Gerold, David Olagnier, Frank Pessler

## Abstract

Nuclear factor erythroid 2–related factor 2 (NRF2) is a master regulator of anti-oxidative and detoxifying cell responses. In addition, it plays important roles in host cell defenses against pathogenic viruses, and small molecules that activate NRF2 signaling can exert potent antiviral effects. We recently found that the NRF2 activators 4-octyl itaconate (4OI), bardoxolone (BARD), and sulforaphane (SFN) interfere with influenza A virus replication by blocking the nuclear export factor exportin 1 (XPO1), which did not require NRF2 signaling. Here, we have assessed their potential to inhibit highly pathogenic (SARS-CoV-2) and seasonal (hCoV-229E) coronaviruses and begun to elucidate the involved mechanisms of action. Using human cell lines and iPSC-derived vascular endothelial cells, we find that NRF2 knock-out or knock-down enhances infection by both viruses, indicating that physiologic NRF2 signaling restricts human coronavirus infection. 4OI, BARD, SFN, as well as the XPO1 blocker Selinexor (SEL), greatly limit infection by both viruses, but in an NRF2-independent manner. Strikingly, the compounds (particularly 4OI) downregulate *ACE2* and *TMPRSS2* mRNA and protein in Calu3 cells, leading to a >10-fold reduction in viral cell entry by 4OI and SEL, as assessed using SARS-CoV-1 and -2 spike protein VSV pseudotypes. A cycloheximide chase experiment revealed that 4OI dramatically reduces ACE2 half-life, which requires the E3 ligases NEDD4L and MCM1, suggesting that 4OI targets ACE2 for destruction by the proteasome. Moreover, 4OI and SEL reduce XPO1 protein levels, and all compounds reduce *XPO1* mRNA levels. Co-incubation experiments of 4OI and the transcription blocker actinomycin D in A549 cells suggest that 4OI acts primarily by interfering with transcription of the *XPO1* gene. XPO1 knock-down markedly reduces 229E replication. All four compounds interfere with 229E infection, but do not alter expression of ANPEP, the cellular receptor for this virus. Their anti-229E efficacy depends on expression of XPO1 in host cells in the order of SEL (most dependent) >4OI >SFN >BARD (least dependent), suggesting that especially BARD interferes with 229E infectivity via yet another, unknown, target. Taken together, these results suggest that “NRF2 activators” act as potent antivirals against human coronaviruses by targeting diverse host factors which are critical for viral infectivity.

**Author summary:** Host-directed antiviral compounds act by a variety of mechanisms. For instance, they stimulate cellular antiviral immune responses and target host cell factors which are required for the viral life cycle. Pharmacologic activation of the NRF2 signaling pathway is a particularly attractive antiviral strategy, as this pathway restricts replication of a variety of viruses and also protects cells from excessive inflammation and oxidative stress resulting from accumulation of reactive oxygen species. In our previous study of the NRF2 activators bardoxolone, sulforaphane, and 4-octyl itaconate as host-directed treatments for influenza A virus infection, we found that these compounds interfered with replication of the virus. Unexpectedly, this antiviral activity was completely independent of NRF2 signaling, but resulted from blocking the nuclear export factor XPO1. In the present study, we find that these compounds limit infection by SARS-CoV-2 and hCoV-229E and that, again, this antiviral effect is NRF-independent. Instead, it depends to a large extent on downregulating ACE2 and TMPRSS2 (the major host cell receptors for SARS-CoV-1 and 2) and blocking/downregulating XPO1. Our results underscore the potential of “NRF2 activators” as adjunct treatments for viral infections, as they protect the host by anti-oxidative, anti-inflammatory, and cytoprotective mechanisms and also interfere with diverse host factors required for the viral life cycle.

## Introduction

Highly pathogenic human coronaviruses (SARS-CoV-1, SARS-CoV-2, MERS) can cause severe infection characterized by hyperinflammation and end-organ damage such as acute respiratory distress syndrome and multi-organ failure, particularly in individuals with poor pre-existing immunity or medical risk factors such as obesity, diabetes, or cardiovascular disease [1]. So called seasonal, low pathogenic coronaviruses (e.g., hCoV-229E, -OC43, -NL63, -HKU1) usually cause mild upper respiratory infections but they can, nonetheless, cause severe disease in immunocompromised individuals. Direct-acting antivirals such as remdesivir and nirmatrelvir/ritonavir have been approved for the treatment of SARS-CoV-2 infection [2] (update accessed Dec. 4, 2024). However, considering the strong contribution of dysfunctional inflammatory responses to organ pathology, adjunct therapy with corticosteroids, IL-6 blockade (tocilizumab), and/or the JAK1/2 inhibitor baricitinib is recommended by the World Health Organization [2] (update accessed Dec. 4, 2024). In addition to direct-acting antivirals and immunomodulatory adjunct treatments, host-directed antivirals would constitute a third line in the armamentarium against coronaviruses. Host-directed antivirals are compounds that stimulate endogenous antiviral mechanisms or interfere with host cell functions that are required for viral infectivity. The KEAP1/NRF2 signaling pathway activates cytoprotective and anti-inflammatory responses in host cells, but also has antiviral functions in itself [3–5]. Activators of this pathway, therefore, constitute promising host-directed interventions for acute viral infections. Indeed, antiviral effects of NRF2 agonists have been demonstrated against a broad spectrum of human pathogenic viruses including SARS-CoV-2 [6], HSV [6], and influenza A [5, 7, 8]. However, in most cases it is not known whether the antiviral effects truly depend on induction of NRF2 signaling or whether other targets contribute. Indeed, in our recent study of NRF2 activators as anti-influenza agents we found that the antiviral effect of the “classic” NRF2 activators bardoxolone-methyl (BARD), sulforaphane (SFN), and 4-octyl itaconate (4OI) did not require functional NRF2-signaling, but was mediated largely by inhibiting nuclear export of viral ribonucleoprotein by the nuclear export factor exportin-1 (XPO1, also known as chromosome region maintenance 1, CRM1) [5]. We also proposed a structural explanation why the compounds recognize such diverse targets: they can bind covalently to the functionally critical cysteine 528 of XPO1 because it is displayed in a structural context very similar to cysteine 151 in their canonical binding site on KEAP1, the major cellular inhibitor of NRF2 [5]. Interestingly, pharmacologic XPO1 inhibition also reduces infectivity of SARS-CoV-2 *in vitro* and *in vivo* even though coronaviruses replicate in the cytoplasm, suggesting that XPO1 mediates shuttling of viral proteins or nuclear export of host cell factors required for optimal infectivity [9, 10].

In the current study, we aimed to assess the relative contributions of activating NRF2 signaling and inhibiting XPO1-mediated nuclear export to anti-coronavirus activity. Specifically, we evaluated the ability of these three NRF2 activators and the selective inhibitor of nuclear export selinexor (SEL) to reduce infectivity of SARS-CoV-2 and the low-pathogenic human coronavirus (HCoV)-229E and tested whether the observed antiviral effects indeed depended on the presumed targets, i.e. NRF2 and XPO1. We find that all compounds exert substantial anti-coronavirus effects, but that these are independent of NRF2 signaling and are in part due to targeting XPO1 and, in the case of SARS-CoV-1 and -2, their cellular receptors ACE2 and TMPRSS2.

## Results

### Inhibition of SARS-CoV-2 is NRF2 independent

We first compared the ability of the NRF2 activators (BARD, SFN, 4OI) to inhibit SARS-COV-2 infection in Calu3 cells 48 h post infection (p.i.). LDH release assay demonstrated that the compounds were non-toxic to Calu3 cells after 48 h of incubation (**Figure S1**). All three compounds significantly reduced viral copy number in cell supernatants and lysates (**Figure 1A,B**). The antiviral effect of 4OI was also verified by immunoblot. 4OI treatment essentially prevented accumulation of SARS-CoV-2 spike and nucleocapsid protein and led to higher levels of the NRF2-inducible proteins AKRB1 and NQO1 independent of SARS-CoV-2 infection, demonstrating its ability to activate NRF2 signaling in this infection model (**Figure 1C**). Since 4OI blocks XPO1 and XPO1 inhibition can lead to reduced XPO1 protein levels [11], we also measured XPO1 expression by immunoblot. Indeed, XPO1 levels were markedly lower under 4OI treatment (**Figure 1C**), which agreed well with the findings by Ribo-Molina et al. that 4OI treatment reduced XPO1 levels in MDCK and Jurkat cells [12]. 4OI efficiently restricted infectivity of SARS-CoV-2 isolates belonging to the α, β, and δ clades in Calu3 cells independent of their ability to replicate in untreated cells (**Figure 1D**). Using 4OI (which had exerted the strongest anti-SARS-CoV-2 effect) as example, we then tested to what extent the antiviral effect of an NRF2 activator actually depends on NRF2 signaling. The CRISPR KO of NRF2 in Calu3 cells led to increased SARS-CoV-2 RNA levels, confirming the antiviral role of endogenous NRF2 signaling (**Figure 1E**). However, 4OI reduced viral copy numbers to the same level in the NRF2 KO cells as in cells electroporated with AAVS1 sequence as negative control. Likewise, reduction of SARS-CoV-2 spike protein by 4OI was only minimally affected by siRNA-mediated knockdown of NRF2 (**Figure 1F**). Taken together, the above results suggest that the antiviral effect of at least 4OI applied to SARS-CoV-2 isolates from different clades and was largely independent of NRF2 signaling. Considering the observed downregulation of XPO1 by 4OI, we then compared the XPO1 inhibitor SEL against the NRF2 activators. SEL did reduce SARS-CoV-2 RNA levels more than 50%, but it was clearly less effective than 4OI and BARD (**Figure 1G**). These results suggest that another target, besides XPO1, contributes to the anti-SARS-CoV-2 effects of at least 4OI and BARD.

**Figure 1.**
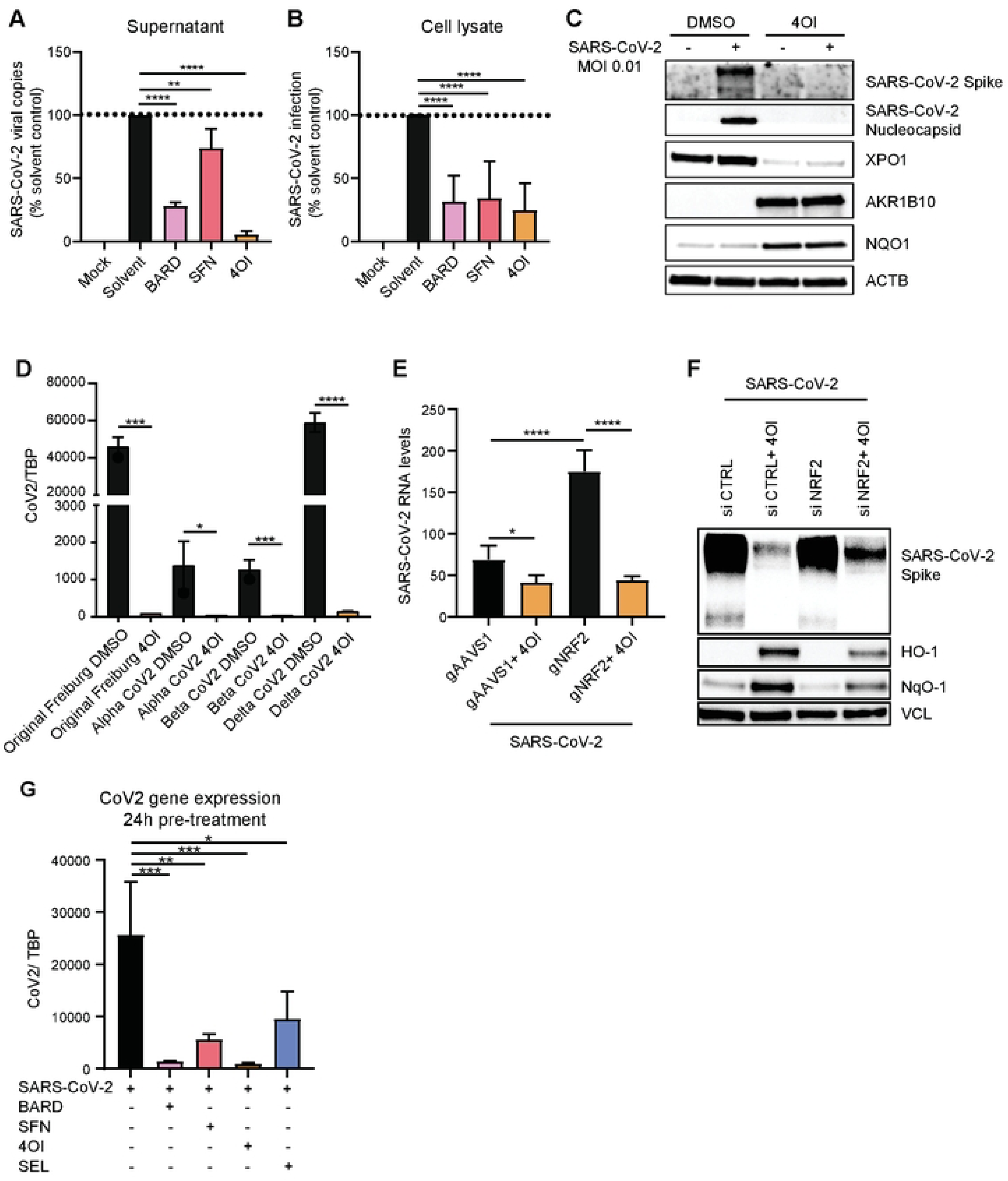
NRF2 activators (4OI, BARD, SFN) and SEL reduce SARS-CoV-2 infectivity in Calu3 cells. Experiments for panels A and B were performed in the Gerold lab (Hannover, Germany) and for panels C-G in the Olagnier lab (Aarhus, Denmark). LDH release assay demonstrated that the compounds were non-toxic to Calu3 cells after 48 h of incubation (**Figure S1**). **A,B**. Calu3 cells were pretreated with the compounds (BARD, 0.1 µM; SFN, 10 µM; 4OI, 100 µM) for 24 h, inoculated with SARS-CoV-2/München-1.2/2020/984,p3 (MOI = 0.005) in presence of the compounds for 4 h, followed by removing the viral inoculum and adding fresh medium containing the respective compounds and controls. 48 h p.i., viral genome copies were determined by RT-qPCR. **A.** supernatants and **B.** cell lysates. **C-G**. Calu3 cells were pretreated with the indicated compounds for 48 h (C-F) or 24 h (G), infected with SARS-CoV-2 Wuhan-like early European B.1 lineage (FR-4286) (MOI = 0.01) for 1 h, followed by removal of the inoculum and incubation in fresh medium containing the compounds. Target gene expression and protein levels were measured 24 h p.i. **C.** Reduction of SARS-CoV-2 spike and nucleocapsid proteins, and XPO1 protein expression, but increase in AKR1B10 and NQO1 levels by 4OI (125 µM) (immunoblot with β-actin as internal reference). **D.** Marked reduction of viral RNA of diverse SARS-CoV-2 variants of concern by 4OI (125 µM). **E,F.** NRF2 independence of the anti-SARS-CoV-2 effect of 4OI. Control (transfected with control siRNA) or NRF2 knock-down Calu3 cells (transfected with specific anti-NRF2 siRNA) were infected with SARS-CoV-2 (MOI 0.01) and treated with 4OI (125 µM) or buffer only. **E**. SARS-CoV-2 RNA (RT-qPCR, cell lysates). **F.** SARS-CoV-2 spike protein (immunoblot with vinculin as internal reference, cell lysates). *n*=3, means ±SEM. One-way ANOVA with Tukey’s post-hoc test. * ≤0.05, ** ≤0.01, *** ≤0.001, **** ≤0.0001.

### Downregulation of ACE2, TMPRSS2, and XPO1

SARS-CoV-2 infection strongly induced expression of interferon (IFN) regulated genes *IFIT1* and *CXCL10* (**Figure 2A,B)**. Consistent with their antiviral effects, all compounds significantly reduced expression of both mRNAs, whereby 4OI led to the greatest reduction, possibly due to a combination of stronger antiviral and anti-IFN effects. We then searched for cellular antiviral targets of the compounds. Motivated by a pilot experiment which showed that BARD interfered with cell entry of vesicular stomatitis virus (VSV) pseudotyped with SARS-CoV-2 spike protein (**Figure S2A**), we measured expression of angiotensin-converting enzyme 2 (ACE2, the main cellular receptor for SARS-CoV-2), transmembrane serine protease 2 (TMPRSS2, the functional co-receptor of SARS-CoV-2), and XPO1. The three compounds reduced levels of all three mRNAs in the order of efficiency 4OI>SFN=BARD (**Figure 2C-E**). Immunoblotting verified that adding 4OI or SEL to uninfected Calu3 cells resulted in rapid reduction of ACE2 and TMPRSS2 protein levels, resulting in complete loss by 48 h (**Figure 2F**), whereas ACE2 levels remained unchanged when the cells were cultured for 72 h in medium not containing the compounds (**Figure S2B**). We then performed a chase experiment with cycloheximide (CHX, which blocks translation) to test whether 4OI affects half-life of ACE2 protein. Addition of CHX to medium resulted in a gradual reduction of ACE2 levels over 12 h (**Figure 2G**), whereas adding CHX and 4OI together led to complete loss of ACE2 protein within 6 h. Thus, 4OI markedly reduced the half-life of ACE2 protein. ACE2 levels can be regulated by ubiquitination, leading to degradation by the proteasome [13]. Surprisingly, the proteasome inhibitor MG132 led to even more rapid loss of ACE2 when given alone and did not reduce ACE2 reduction by 4OI (**Figure S2C,D**), suggesting that it may induce ACE2 degradation by an independent mechanism. For instance, MG132 has been shown to induce autophagy [14]. We then tested the hypothesis that 4OI induces activity of the e3 ligases that contribute the most to ubiquitination, and thus proteasomal degradation, of ACE2. A network analysis of predicted E3 ligase targets of ACE2 revealed multiple interactions, with strongest functional relationships with MDM2 and NEDD4L (**Figure S3**). Indeed, knocking down *NEDD4L* mRNA with siRNA nearly completely abrogated the ACE2-destroying capacity of 4OI (**Figure 2H**). siRNA knock-down of *MDM2* attenuated ACE2 destruction by 4OI was well, but the effect was less pronounced (**Figure 2J**). Of note, knock-down of either ligase also mitigated the reduction of *ACE2* mRNA by 4OI and SEL (**Figure 2I, K**).

**Figure 2.**
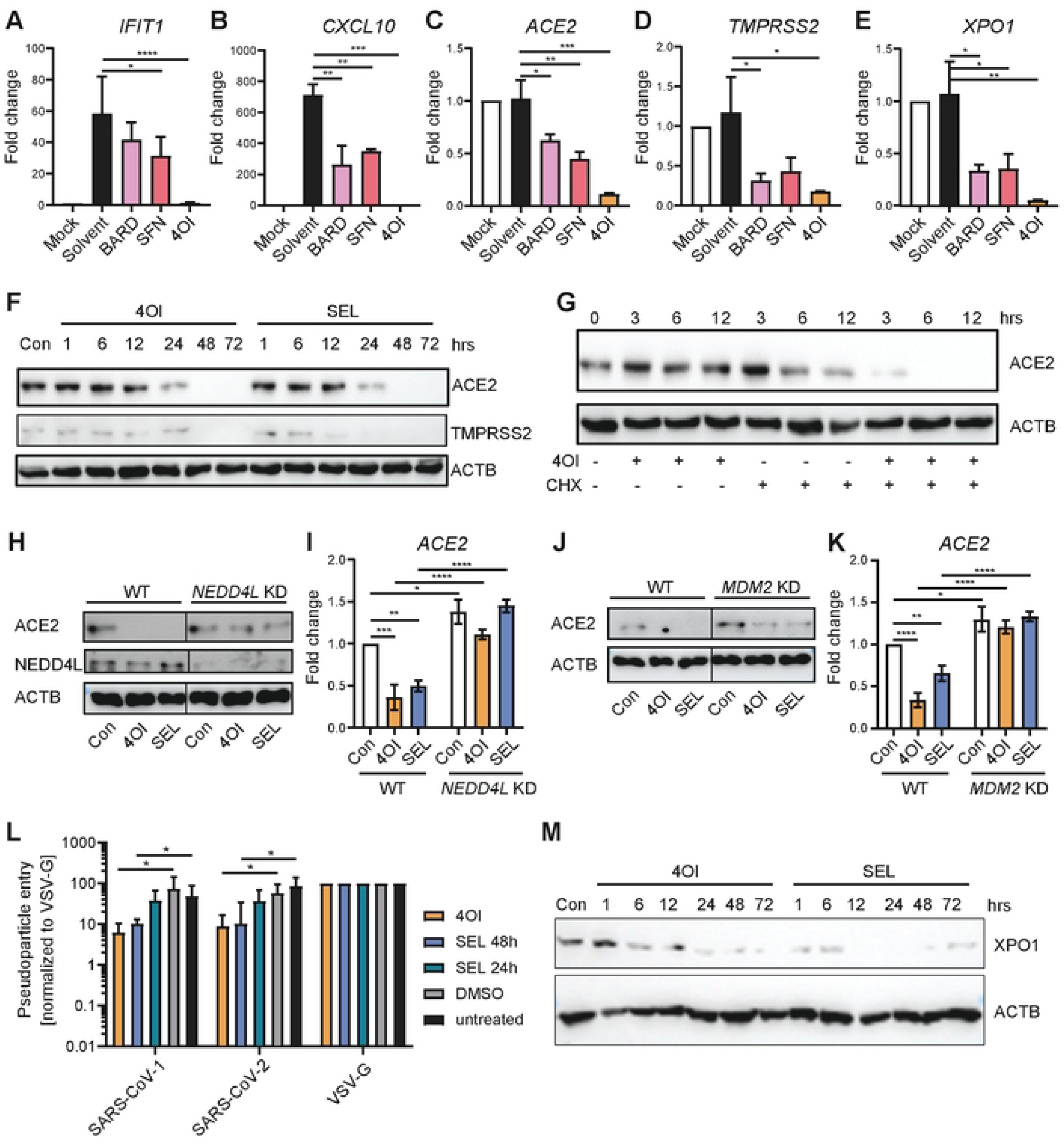
4OI reduces ACE2 and XPO1 levels and interferes with SARS-CoV-1 and SARS-CoV-2 cell entry. **A-E.** Reduction of *IFIT1*, *CXCL10*, *ACE2*, *TMPRSS2*, and *XPO1* mRNA. RT-qPCR analysis of RNA from the experiment shown in Figure 1A**,B**. **F.** 4OI and SEL reduce ACE2 and TMPRSS2 protein levels. Uninfected Calu3 cells were grown in medium containing 4OI (100 µM) or SEL and cellular ACE2 and TMPRSS2 levels were measured by immunoblot after 1 – 72 h. **G**. 4OI reduces half-life of ACE2. Uninfected Calu3 cells were grown in medium containing 4OI (100 µM) with our without cycloheximide (CHX, 50 µg/ml) and cellular ACE2 levels were measured by immunoblot after 1 – 12 h. **H, I.** NEDD4L knock-down attenuates the ACE2-destroying capacity of 4OI at the protein (H) and the mRNA level (I). **J, K.** MDM2 knock-down attenuates the ACE2-destroying capacity of 4OI at the protein (J) and the mRNA level (K). **L.** 4OI and SEL interfere with cell entry of SARS-CoV-1 and -2. Calu3 cells were pre-incubated with 4OI (48 h) or SEL (24 h or 48 h) and inoculated with luciferase-expressing VSV particles pseudotyped with SARS-CoV-1 or -2 spike protein or VSV-G protein. Cell entry was assessed by luciferase activity 16 h after inoculation. SARS-CoV pseudotype luciferase signal was normalized against the signal obtained with VSV-G pseudotypes, which was set as 100%. Pooled analysis of 6 independent experiments with *n*=3 replicates each. **M.** 4OI and SEL reduce XPO1 protein levels. Uninfected Calu3 cells were grown in medium containing 4OI or SEL, and XPO1 levels were measured by immunoblot 1 - 72 h after addition of the compounds. A-E and H: One-way ANOVA with Tukey’s post-hoc test. * ≤0.05, ** ≤0.01, *** ≤0.001, **** ≤0.0001.

In cell entry assays with a VSV vector pseudotyped with SARS-CoV-2 or SARS-CoV-1 spike protein (or VSV-G protein as positive control for ACE2-independent cell entry), 4OI and SEL (which enhances nuclear localization of ACE2 [9]) markedly reduced entry of SARS-CoV-2 as well as SARS-CoV-1, which also uses ACE2 as main cellular receptor (**Figure 2L**). Thus, both 4OI and SEL interfered with SARS-CoV-2 infectivity at the level of cell entry mediated by spike protein. Considering that 4OI treatment reduced XPO1 protein in uninfected and infected Calu3 cells (**Figure 1C)** and *XPO1* mRNA in infected Calu3 cells **(Figure 2E**), we then compared the ability of 4OI and SEL to reduce XPO1 protein levels. XPO1 levels decreased greatly in the presence of either compound, with SEL exerting a somewhat stronger effect (**Figure 2M**). Taken together, these results suggest that downregulation of ACE2, TMPRSS2, and XPO1 is a major component of the anti-SARS-CoV-2 effect of 4OI and SEL, and, to a lesser extent, also BARD and SFN.

### NRF2-independent inhibition of seasonal coronavirus hCoV-229E

This low-pathogenic coronavirus can cause severe disease in immunocompromised individuals. It shares cardinal features of viral replication with SARS-CoV-2, but can be studied under biosafety level 2 regulations. We therefore focused further studies on this pathogen. As opposed to SARS-CoV-2, hCoV-229E can productively infect microvascular endothelial cells (ECs) [15]. Indeed, a *Renilla* luciferase-encoding hCoV-229E strain (here referred to as 229E-luc) replicated efficiently in hiPSC-derived ECs, as evidenced by vigorous luciferase activity and high levels of RNA encoding the viral M protein (**Figure 3A,B**). As observed with SARS-CoV-2 in Calu3 cells (see **Figure 1E**), deleting the *NFE2L2* gene (which encodes NRF2) led to significantly increased viral replication, lending further support to its importance in restricting coronavirus infection in human cells. However, the three NRF2 activators (as well as SEL) prevented viral replication nearly completely in both the WT and *NRF2^−/-^* ECs, indicating that their antiviral effect does not depend on NRF2 signaling. Of note, in contrast to *ACE2* and *TMPRSS2* mRNA in Calu3 cells (see **Figure 2C,D**), the compounds did not affect expression of mRNA encoding ANPEP, the major cellular receptor for hCoV-229E [16] (**Figure 3C**). However, as in Calu3 cells, the three NRF2 activators reduced *XPO1* mRNA levels, with 4OI causing the greatest reduction (**Figure 3D**). A similarly pronounced reduction was observed by SEL (**Figure 3D)** and the naturally occurring XPO1 inhibitor leptomycin B, (**Figure S4**), suggesting that the reduction in *XPO1* mRNA was mediated, at least indirectly, by binding of the compounds to XPO1 protein. Co-treatment with SEL and 4OI, BARD, or SFN resulted in approximately the same reduction of *XPO1* mRNA levels than treatment with SEL alone. *XPO1* mRNA levels were significantly higher in the *NRF2^−/-^* ECs, but the treatments reduced its expression to the same levels as in the treated WT ECs (**Figure 3D, Figure S4**). Thus, downregulation of *XPO1* mRNA by the compounds did not require NRF2 signaling.

**Figure 3.**
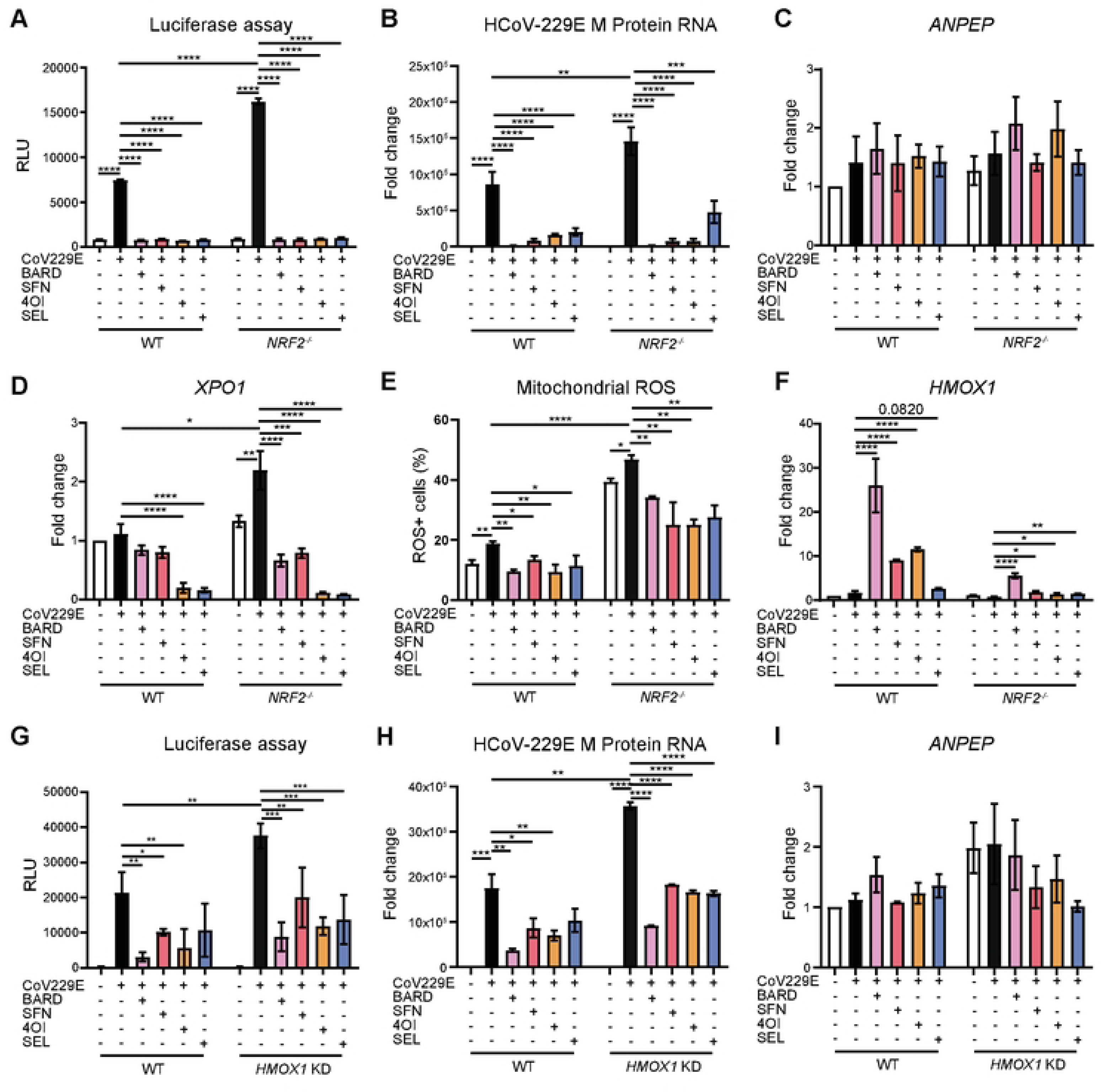
NRF2-independent inhibition of 229E infectivity and downregulation of *XPO1* mRNA. **A-F**. WT and *NRF2^−/-^* human iPSC-derived ECs were infected with the luciferase-labeled strain hCoV-229E-luc (MOI=0.3) for 4 h and then cultured for 48 h in fresh medium containing the compounds. Luciferase activity, viral M protein RNA levels, host mRNA expression, and mitochondrial ROS levels were measured after 48 h. **A,B.** Luciferase activity and viral M protein RNA. **C.** *ANPEP* mRNA. **D.** *XPO1* mRNA. **E.** Mitochondrial ROS (flow cytometry). **F.** *HMOX1* mRNA. **G-I.** Effect of HMOX1 knock-down on 229E-luc infectivity and antiviral activity of the compounds in A549 cells. Infections and treatments were carried out as in A-F, except that WT (transfected with scrambled control siRNA) or siRNA-mediated *HMOX1* knock-down A549 cells were used. **G.** Luciferase activity. **H.** M protein RNA. **I**. *ANPEP* mRNA. One-way ANOVA with Tukey’s post-hoc test. * ≤0.05, ** ≤0.01, *** ≤0.001, **** ≤0.0001.

Levels of reactive oxygen species (ROS) were nearly 3-fold higher in uninfected *NRF2^−/-^* than wild-type ECs, underscoring the anti-oxidative impact of NRF2 signaling (**Figure 3E**). 229E-luc infection led to a modest ROS increase in both *NRF2^−/-^* and wild-type cells, and the four treatments reduced ROS levels in the infected *NRF2^−/-^*cells below the level measured in the uninfected *NRF2^−/-^* cells. However, even though the three NRF2 activators had essentially abolished 229E-luc infection (**Figure 3A-B**), ROS levels remained significantly higher in the treated *NRF2^−/-^* cells than in the treated WT cells, suggesting that the anti-oxidative effects of these compounds depended at least partially on NRF2 signaling. Indeed, the compounds induced the anti-oxidative NRF2 target gene *HMOX1* only weakly in the *NRF2^−/-^*cells (**Figure 3F)**. HMOX1 has also been implicated in antiviral functions of NRF2 [17]. We therefore assessed the impact of siRNA-mediated knock-down of *HMOX1* mRNA on 229E-luc infectivity (**Figure 3G-I**). Considering the susceptibility of these iPSC-derived ECs to technical artifacts from siRNA transfection, we performed this experiment in A549 cells. These cells supported more vigorous replication of the virus (compare the lower Luc signals obtained in ECs, shown in **Figure 3A**), but the compounds were also somewhat less effective. An approx. 80% knock-down of *HMOX1* mRNA was achieved (**Figure S5H**). Luciferase activity and M protein RNA expression were significantly higher in the *HMOX1* knock-down cells, supporting its antiviral role. Percent reduction of both parameters by the three NRF2 activators was slightly lower in the HMOX1 knock-down cells (**Table S1**), indicating that it does contribute to their antiviral effect. SEL was somewhat more effective in the *HMOX1* knock-down cells, suggesting that HMOX1 interferes with its activity. This may relate to the previously described interaction between heme and XPO1-mediated nuclear export [18]. Neither HMOX1 knock-down nor the treatments affected expression of *ANPEP* mRNA (**Figure 3I**).

### XPO1 is required for optimal infectivity of 229E and the antiviral effect of 4OI

Having demonstrated the NRF2-indepence of the compounds’ anti-229E effects, we then assessed the contribution of XPO1 function to 229E infectivity and to their anti-229E activity. A 98% knockdown was achieved with anti-XPO1 siRNA in A549 cells, which was greater than the reduction seen after treatment with SEL (97%), 4OI (88%), Bard (59%), and SFN (59%) (**Figure 4A**). The XPO1 knockdown led to a substantially larger reduction of M protein RNA (∼80%) than of luciferase activity (∼50%), which was comparable in both cases to the reduction by SEL treatment of WT cells (**Figure 4B,C)**. One can thus conclude that XPO1 contributes greatly to, but is not strictly required for, 229E infectivity in these A549 cells. As expected, SEL treatment of XPO1 knockdown cells did not reduce M protein RNA level or luciferase activity much further, confirming that, among the compounds tested herein, the antiviral activity of SEL depends the most on the presence of XPO1. In contrast, a further reduction in these parameters ensued under treatment with BARD and, to a lesser extent, SFN and 4OI, suggesting that especially BARD acts at least partially through an additional target. A comparison of differences in percent reduction in 229E infectivity by the compounds between WT and knock-down cells suggested the following order of XPO1 dependence of the compounds: SEL>4OI>SFN>BARD, which correlates well with their predicted affinities for the XPO1 NES binding site [5]. XPO1 knock-down did not affect expression of *ANPEP* mRNA, but it slightly reduced the ability of BARD to induce *NFE2L2* and *HMOX1* mRNA in 229E-infected cells, the functional relevance of which is uncertain (**Figure 4D-F**).

**Figure 4.**
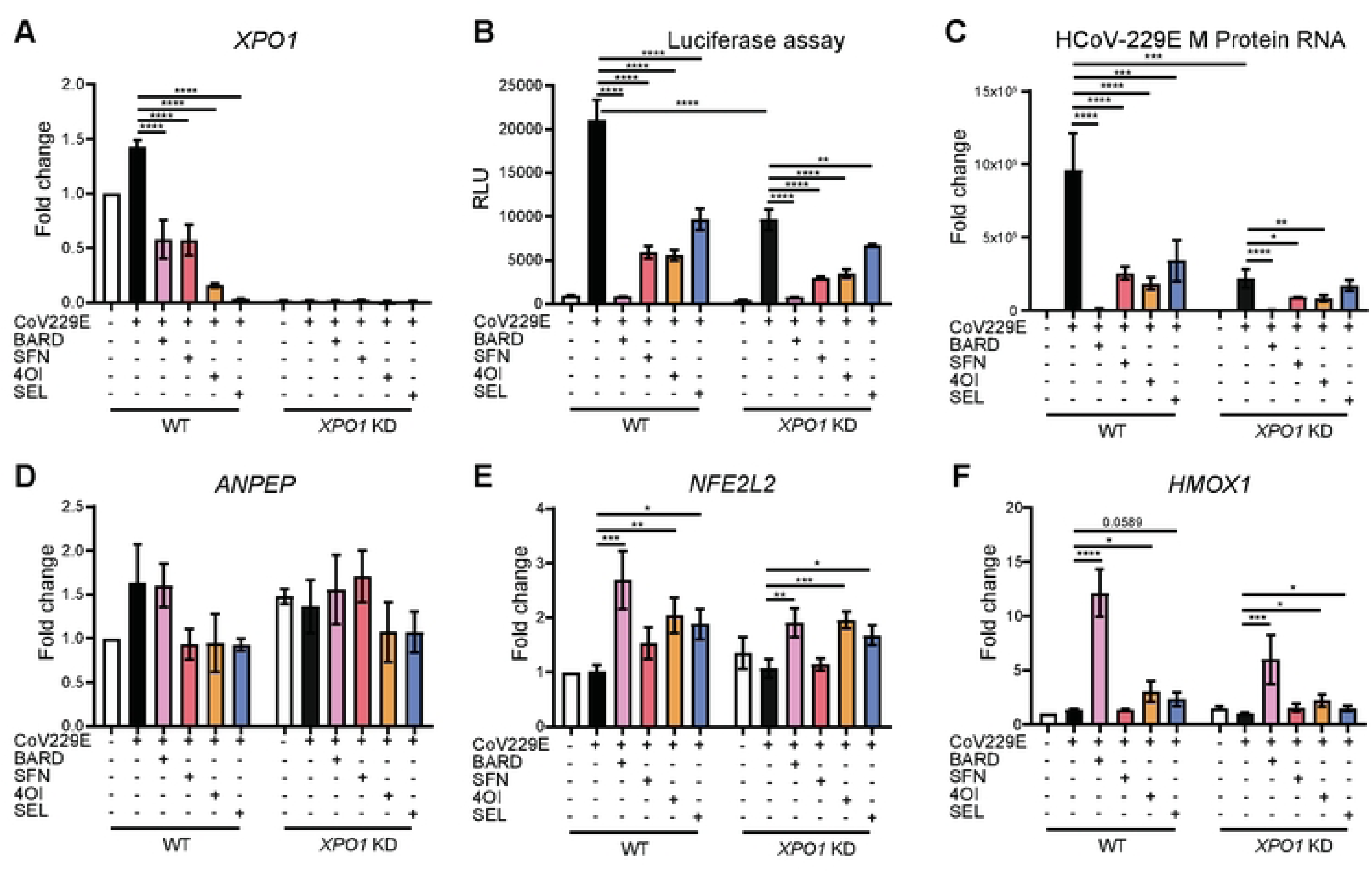
XPO1 knock-down reduces infectivity of hCoV-229E. *XPO1* mRNA was knocked down in A549 cells by siRNA, and infectivity of 229E-luc and antiviral efficacy of the compounds were assessed in WT (transfected with scrambled control siRNA) and XPO1 KD cells following the same infection protocol as in Figure 3. **A.** *XPO1* mRNA (RT-qPCR). **B.** Luciferase activity. **C.** M protein RNA expression. **D.** *ANPEP* mRNA (RT-qPCR). **E.** *NFE2L2* mRNA (RT-qPCR). **F.** *HMOX1* mRNA (RT-qPCR). *n*=3. One-way ANOVA with Tukey’s post-hoc test. * ≤0.05, ** ≤0.01, *** ≤0.001, **** ≤0.0001.

### 4OI reduces *XPO1* mRNA expression predominantly at the level of transcription

We selected *XPO1* mRNA to explore the mechanism of downregulation at the transcriptional level because (as opposed to ACE2) it relates to both SARS-CoV-2 and 229E. siRNA-mediated knock-down of KEAP1 (leading to NRF2 activation) in human cells greatly shortens the half-life of *STING* mRNA, and addition of 4OI downregulates *STING* mRNA as well [19]. However, it is not known whether 4OI acts via the same post-transcriptional mechanism as KEAP1 inactivation. Considering that 4OI blocks nuclear export via XPO1, we performed a pathway analysis using XPO1 cargoes documented in the ValidNESs database [20] (**Figure 5A**). Relevant enriched pathways included cellular functions involving macromolecule catabolism (*autophagy*), RNA turnover (*mRNA surveillance*, *RNA degradation*), but also RNA synthesis (*RNA polymerase*) and its activation (*NF-κB signaling*). Thus, XPO1 inhibition could affect mRNA levels of a given target by multiple mechanisms. We first tested the effect of 4OI on *XPO1* mRNA half-life in uninfected A549 cells in the presence and absence of actinomycin D (ActD), which shuts down cellular transcription and allows measuring the natural decay of a given mRNA. *XPO1* mRNA levels decreased with similar kinetics when 4OI and ActD were applied as single treatments (**Figure 5C**). If 4OI shortens *XPO1* mRNA half-life by a mechanism downstream of ActD, e.g. by increasing mRNA turnover, then the half-life should be shorter when both compounds are added together. Surprisingly, incubating the cells in medium containing both 4OI and ActD actually prolonged the half-life of *XPO1* mRNA (20.5 h vs. 10-12 h). The cytoprotective effects of 4OI are well documented and to a large extent are mediated by induction of NRF2 signaling [21, 22]. We hypothesized that 4OI diminished the effect of ActD via the efflux pump ABCB1, which is encoded by an NRF2 target gene and exports ActD from cells [23]. Indeed, when we knocked down *ABCB1* mRNA with siRNA (**Figure 5B**), the half-life-prolonging effect of 4OI/ActD co-treatment was greatly diminished, whereas the effect of ActD single treatment was enhanced, suggesting that 4OI attenuates the effect of ActD in an ABCB1-dependent manner (**Figure 5D**). The hypothesis that 4OI and ActD act via a similar mechanism is also supported by differences between WT and knock-down cells when 4OI was applied after ActD. When WT cells (in which *ABCB1-*mediated export is expected to result in lower steady-state ActD levels) were treated for 24 h with ActD and then grown in fresh medium containing 4OI, *XPO1* mRNA levels continued to decline (**Figure 5C**, right panel). On the other hand, in *ABCB1* knock-down cells (where ActD levels are expected to be higher and *XPO1* mRNA levels were indeed significantly lower at 24 h) a further reduction by the 4OI-containing medium was not observed (**Figure 5D**, right panel). Taken together, these results suggest that 4OI predominantly interferes with *XPO1* mRNA expression at the same step in gene expression as ActD, i.e. by inhibiting transcription. However, considering the potential confounding interactions between ActD and 4OI we cannot fully exclude a contributing mechanism at the post-transcriptional level. This would also agree with the close association of XPO1 function with RNA turn-over mentioned above (**Figure 5A**).

**Figure 5.**
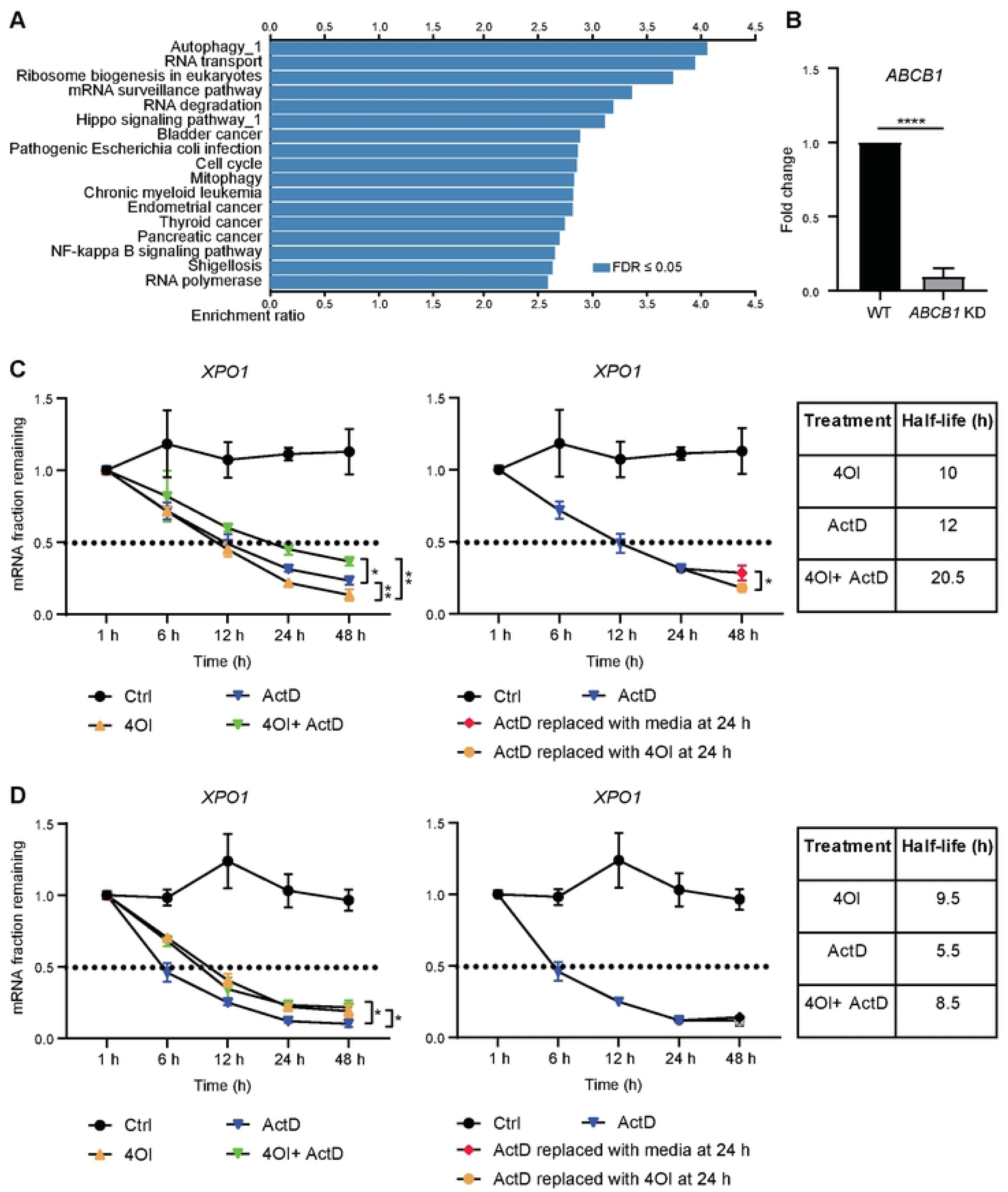
4OI reduces transcription of the *XPO1* gene. **A.** KEGG pathway analysis based on XPO1 cargoes listed in the ValidNESs database [20]. **B.** Efficiency of *ABCB1* mRNA knock-down with siRNA (RT-qPCR). **C.** Comparison of half-life reduction of *XPO1* mRNA by ActD, 4OI, or co-treatment with both. Numerical values for half-life were obtained by extrapolation and are listed in the table next to the graph. **D.** Same experiment as in **C** but using *ABCB1* knock-down (siRNA) A549 cells. Representative of two independent experiments, *n*=3. One-way ANOVA with Tukey’s post-hoc test. * ≤0.05, ** ≤0.01, *** ≤0.001, **** ≤0.0001.

### Changes in host cell transcriptomes due to hCoV-229E infection and treatment with 4OI, BARD, and SEL

We then used RNAseq to characterize the impact of 229E infection on host cell transcriptomes and to test whether the compounds differed in their effects on transcriptomes of infected cells. As expected, 229E infection of A549 cells led to major transcriptional reprogramming, both at 24 and 48 h p.i., which largely normalized upon treatment with all three compounds (**Figure 6A**). For instance, 229E-luc infection led to a major increase in variance compared to mock-infected samples along both PC1 and 2 at 24 h, which increased further by 48 h, whereas the treated samples clustered close to the mock-infected samples at both time points. The Venn diagrams illustrate that there were differentially expressed (DE) genes unique to each compound and shared between two compounds, but that the largest numbers of DE genes (>5000 at both time points) were commonly DE upon treatment with all three compounds (**Figure 6B** (48 h) **Figure S6A** (24 h)). In agreement with this, a gene set enrichment analysis (GSEA) based on KEGG pathways revealed that the three compounds affected very similar functional pathways in infected host cells (**Figure 6C** (48 h), **Figure S6B** (24 h)). Targeted RT-qPCR analysis revealed that induction of the classic IFN-stimulated genes *IFIT1* and *CXCL10* was much lower in 229E infection than in SARS-CoV-2 infection (compare **Figure S5** to **Figure 2A,B**), which agreed with prior knowledge that differential gene expression in 229E-infected human cells does not feature a strong IFN response [24]. Indeed, the RNAseq analysis revealed that 229E infection resulted in minimal regulation of the 72 genes making up the KEGG pathway *Type I interferon response*: there were only 3 DE genes at 24 h p.i. (all up), and 5 at 48 h p.i. (2 up, 3 down) (**Figure S6C, Figure 6D**). Treatment with BARD, 4OI, and SEL all reversed these expression changes. In contrast, infection led to a major downregulation of promitotic genes and upregulation of key antiproliferative genes such as *TP53* and *MYC*, which was remarkably reversed by treatment with all three compounds (**Figure 6E, Figure S6D**). The weak IFN response was confirmed by the absence of IFN-related KEGG pathways in the GSEA comparing infected vs. uninfected cells (**Figure 7** (48 h)**, Figure S7** (24 h)). This was not due to an inability of A549 cells to support IFN responses, as infection with IAV leads to brisk IFN-responses in this cell type [25]. Rather, the GSEA revealed an overall cytostatic effect of the infection, as evidenced by significant depletion of pathways relating to cell cycle, nucleotide biosynthesis, and metabolism, which was reversed by treatment with the three compounds. The observation that the compounds normalized cell transcriptomes with such similar patterns suggests that the RNAseq analysis mostly reflects downstream effects of diminished viral replication and not specific modes of action of the compounds.

**Figure 6.**
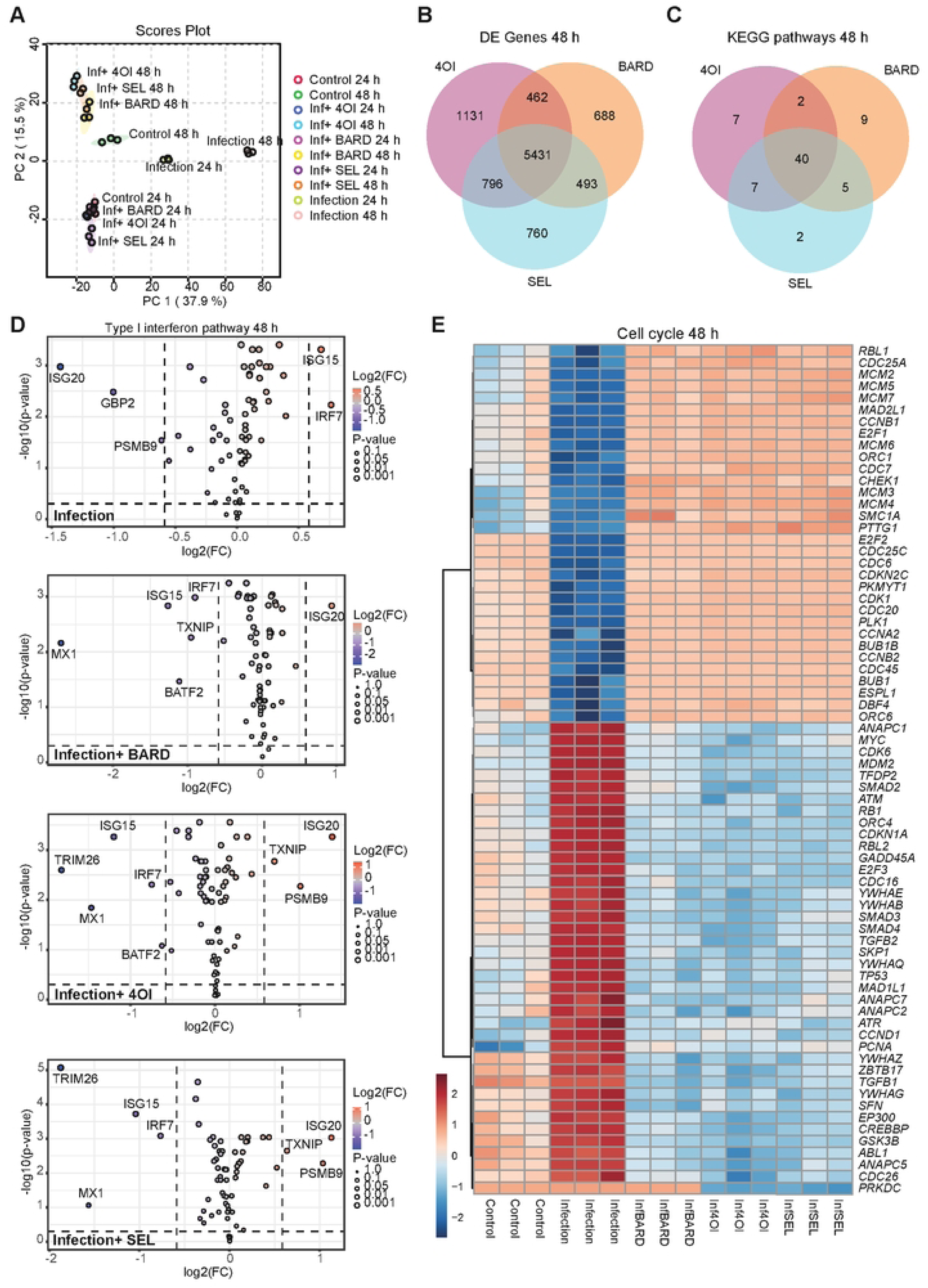
Effect of BARD, 4OI, and SEL on transcriptomes of 229E-infected A549 cells. A549 cells were infected with 229E-luc, treated with the compounds as indicated, and cell transcriptomes were determined by RNAseq 24 and 48 h p.i. **A.** PCA showing centroids of values obtained from the 3 samples in each group. Controls = mock-infected samples. **B,C.** Venn diagrams indicating shared and unique DE genes (**B**) and KEGG pathways (**C**) 48 h p.i. **D.** DE of the genes contained in KEGG pathway *Type I interferon pathway* due to 229-luc infection without treatment and under treatment with BARD, 4OI, and SEL. **E.** Hierarchical clustering analysis (y-axis) of 72 of the 113 genes contained in KEGG pathway *Cell cycle*, which were selected by highest |log2FC| values. The color scale on the lower left indicates *Z*-score.

**Figure 7.**
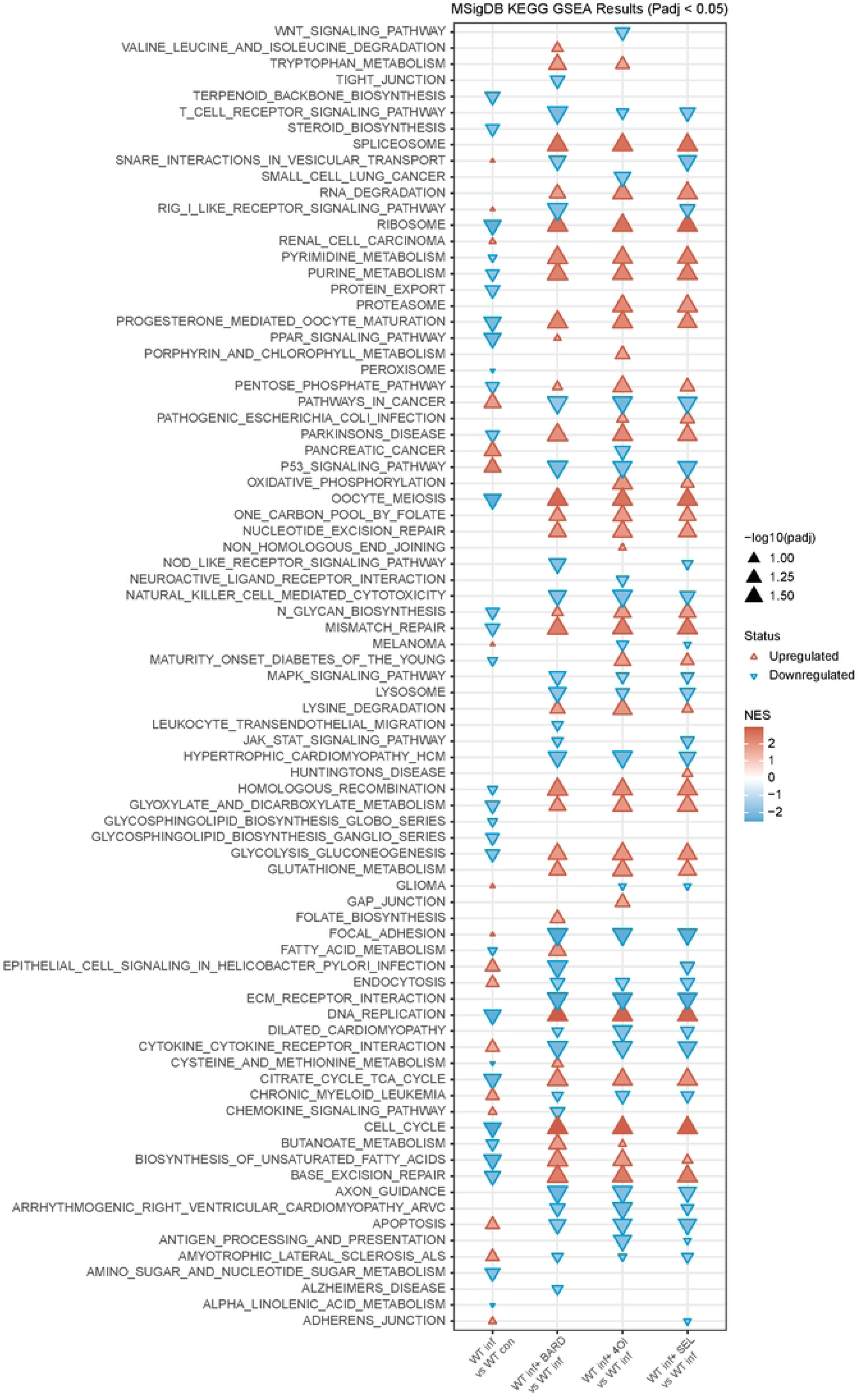
hCoV-229E infection represses signaling pathways associated with cell proliferation and metabolism, which is reversed by treatment with BARD, 4OI, and SEL. GSEA (KEGG pathways) based on the RNAseq data 48 h p.i. used for Figure 6. KEGG pathways were selected by pAdj <0.05 and are listed in reverse alphabetical order top to bottom. NES, normalized enrichment ratio.

## Discussion

In this study of host-directed antivirals against high and low pathogenic human coronaviruses, we find that small molecules which activate the antiviral NRF2 signaling pathway greatly interfere with viral infectivity, but in an NRF2-independent manner. These results agree well with our previous investigations into the mode of action of the same compounds against influenza A virus, where we found that the antiviral activity did not depend on NRF2 signaling but was largely mediated by blocking the nuclear export factor XPO1 [5].

### Antiviral mode of action of BARD

Antiviral efficacy of 4OI and BARD against SARS-CoV-2 has previously been demonstrated in cellular models [6, 26]. Sun et al. suggested that BARD may have direct anti-SARS-CoV-2 activity, presumably by binding to its 3C-like protease [26]. However, the half-maximal effective concentration (EC_50_) to inhibit SARS-CoV-2 replication in cellular infection models was substantially lower than the half-maximal inhibitory concentration (IC_50_) to inhibit the enzyme in a cell-free assay, suggesting the presence of an additional, likely cellular, target. We find that BARD interferes with both SARS-CoV-2 and hCoV-229E. The latter expresses a 3c-like protease with a similar structure, but it is not known whether it can be inhibited by BARD. BARD did reduce expression of ACE2 and TMPRSS2, but the effect was relatively modest, and it inhibited SARS-CoV-2 spike protein-mediated cell entry less than 4OI. In addition, its XPO1 blocking activity is markedly lower than that of 4OI or SEL [5]. The anti-229E activity of BARD did not require NRF2, was less XPO1-dependent, and it did not affect ANPEP expression, suggesting either a direct antiviral effect or the presence of another cellular target. It has been suggested that BARD inhibits the PI3K/Akt/mTOR and p38 MAPK signaling pathways [27]. These pathways are induced by a variety of viral pathogens, including coronaviruses [28, 29] and influenza A virus [7, 30] and are proposed targets for host-directed antivirals. However, inhibiting these pathways by BARD induced cell cycle arrest in K562 cells, i.e., the opposite of what was observed by us in 229E-infected A549 cells. Moreover, the GSEA analysis did not reveal regulation of these pathways by 229E infection or BARD treatment. Thus, a cellular target(s) which mediate(s) the presently unexplained portion of BARD’s anti-CoV effect remains to be identified.

### Antiviral mode of action of 4OI and SEL

Olagnier et al. previously showed that 4OI treatment of SARS-CoV-2 infected cells relieved suppression of NRF2 signaling by the virus, but they did not provide experimental proof that this reconstitution of NRF2 signaling also mediated 4OI’s antiviral effect. The present study now provides evidence that the anti-SARS-CoV-2 effect of 4OI (as well as SEL) is mediated to a large extent by inducing degradation of the major cellular receptor ACE2 and the co-receptor TMPRSS2, downregulating mRNA encoding ACE2 and TMPRSS2, and downregulating XPO1 mRNA and protein. Cellular ACE2 levels are regulated by ubiquitination [31], including by the E3 ligases NEDD4L [32] and MDM2 [33]. Indeed, both E3 ligases were necessary for 4OI to effect destruction of ACE2. In agreement with this, genetic ablation of ACOD1 (the enzyme that synthesizes itaconic acid in activated myeloid cells) leads to major alterations in cellular ubiquitination patters [34]. Of note, SEL induces ubiquitination and subsequent proteasomal degradation of XPO1 [35]. Considering that 4OI and SEL reduced ACE2 levels with similar kinetics (**Figure 2F**), we favor the hypothesis that the two compounds induce ACE2 destruction by similar proteasome-mediated mechanisms. A proteasome-mediated degradation of an activating transcription factor(s) would also explain the observed downregulation of *ACE2*, *TMPRSS2*, and *XPO1* mRNAs. Indeed, STAT3 is a major activator of the ACE2 promoter whose activity can be regulated by proteasomal degradation [36].

In our previous study of the same NRF2 activators and SEL as host-directed interventions against influenza A virus in human cells, we found that the major mode of action of 4OI was to inhibit export of viral ribonucleoprotein by covalently blocking the NES-binding site of XPO1 [5], and this finding was subsequently corroborated by [12]. In addition, 4OI treatment potentiated nuclear localization of p53 in IAV-infected cells [5]. The present study not only revealed the importance of XPO1 for infectivity of both SARS-CoV-2 and 229E, but it also suggests that XPO1 mediates at least part of the anti-CoV activity of 4OI. This appeared counterintuitive because, as opposed to influenza viruses, coronaviruses complete the major phases of their life cycle in the cytoplasm. However, studies of SARS-CoV-1 have shown that viral proteins shuttle between cytoplasm and nucleus (summarized in [9]). Most importantly, the pronounced interference of SEL with cell entry of both SARS-CoV-1 and -2 spike protein pseudotypes clearly demonstrates that blocking XPO1 function can interfere with viral infection, whereby XPO1 inhibition leads to a secondary loss of host cell factors critical for the viral life cycle, in this case by reducing receptor density on target cells. This effect likely also applies to other cell types that express ACE2 and support SARS-CoV-2 infection, such as intestinal epithelial cells, as XPO1 is expressed ubiquitously.

Olagnier et al. had shown that activating NRF2 signaling by knocking down KEAP1 expression led to loss of *STING* mRNA by dramatically shortening its half-life, i.e., by a post-transcriptional mechanism [19]. In their work, addition of 4OI led to loss of *STING* mRNA in an NRF2-dependent manner, but it was not tested whether 4OI acted via the same post-transcriptional mechanism. In contrast, our ActD experiment (**Figure 5**) suggests that 4OI largely acted by interfering with *XPO1* transcription. It has been reported that pharmacologically blocking XPO1 leads to loss of XPO1 protein [37, 38]. *XPO1* mRNA levels have been measured under treatment with KPT-335 (verdinexor) [37] and KPT-185 [38]. Intriguingly, expression was found to be increased, whereas in our study treating A549 cells with SEL (KPT-330) clearly reduced *XPO1* mRNA levels. Thus, the mechanism underlying the loss of XPO1 protein may differ depending on factors such as the pharmacologic agent and cell type studied. In any case, 4OI downregulated XPO1 more efficiently than SEL, suggesting that it interferes with XPO1 expression also through an XPO1-independent mechanism. Indeed, recent work revealed inhibition of TET2 dioxygenase (an enzyme which tends to activate gene transcription by reversing promoter methylation) as a major mechanism of the anti-inflammatory effect of 4OI and endogenous itaconic acid. Other plausible mechanisms include interference with NF-κB activity, for instance by preventing export of IκB-α from the nucleus, as has been described for SEL in breast cancer cells [39]. 4OI’s anti-229E activity appeared to be partially independent of XPO1, and it did not affect expression of its cellular receptor, ANPEP (**Figure 4**), suggesting the presence of other targets. Further research is required to gauge the importance of XPO1-independent mechanisms to 4OI’s anti-CoV activity.

### Importance of endogenous NRF2 signaling

The NRF2 signaling pathway has been extensively implicated in host defenses against pathogenic viruses, and HMOX1 is considered a downstream mediator of this antiviral effect [17]. Our results underscore the strong anti-oxidative function of endogenous NRF2 signaling even in uninfected cells. Moreover, they confirm its antiviral capacity, as replication of both SARS-CoV-2 and 229E increased upon inactivation of the *NFE2L2* gene (**Figure 1E**, **Figure 3A,B**), which was paralleled by a decrease in *HMOX1* expression. Olagnier et al. showed that knocking down *HMOX1* expression did not affect SARS-CoV-2 infection or replication in African Green Monkey Vero cells [6]. In the present study, we tested the effect of *HMOX1* knock-down on 229E infection, which revealed that HMOX1 contributed to restriction of 229E and to the anti-229E effect of the compounds, albeit its contribution to the latter was relatively minor. The activity of HMOX1 against human coronaviruses may thus vary depending on the virus and/or cell type and host species.

In summary, we find that the anti-CoV activity of the NRF2-activating compounds 4OI, BARD, and SFN (as well as the XPO1 inhibitor SEL, which is also predicted to bind to the NRF2 inhibitor KEAP1 [5]) does not require an intact NRF2 signaling pathway and that a good part of the mode of action of 4OI is mediated by downregulating ACE2 and TMPRSS2 and blocking/downregulating XPO1. The latter agrees well with our previous ligand-target modelling results, which suggested that these NRF2 activators and SEL can recognize each other’s canonical binding sites (KEAP1 and XPO1) due to the presence of cysteines which are displayed in similar environments that are conducive to formation of covalent bonds with electrophilic double bonds in the compounds [5]. The higher affinity of 4OI, compared to BARD and SFN, for XPO1 was explained by additional hydrophobic interactions of its 4-octyl tail with the hydrophobic cleft of XPO1. Considering the ability of electrophilic compounds to interact with a multitude of nucleophilic targets in the cell, their complete mode of antiviral action likely results from the sum of diverse additive, counteractive, and synergistic interactions which may vary according to cell type, activation status, and life cycle of the virus. Considering that they integrate antiviral, anti-inflammatory, and cytoprotective properties, electrophilic NRF2-activating small molecules like 4OI, SFN, and BARD certainly constitute promising starting points for the development of host-directed antivirals.

## Methods and Materials

### Compounds

Bardoxolone methyl (BARD; 2-cyano-3,12-dioxo-oleana-1,9(11)-dien-28-oic acid methyl ester, Hölzel/MedChem Express, Sollentuna, Sweden, HY-13324-50mg), 4-octyl itaconate (4OI; Biomol Cayman chemical, Hamburg, Germany, Cay25374-25), sulforaphane (SFN; (*R*)-1-isothiocyanato-4-(methylsulfinyl)-butane; Santa Cruz, Heidelberg, Germany, sc-203099), selinexor (SEL; selective inhibitor of nuclear export KPT-330, Tebu-Bio, Offenbach, Germany, 10-4011-25 mg). We used previously documented nontoxic concentrations: 4OI 100 µM [7], SEL 1 µM [40], BARD 0.1 µM [41], SFN 10 µM [42]. The lack of toxicity of the compounds at these concentrations on Calu3 cells was additionally verified by LDH release assay (**Figure S1**). Leptomycin B (LMB, 431050-5UG, Sigma Aldrich) was applied at a concentration of 20 nM, and MG132 (S2619, Hölzel Diagnostica) at 10 µM.

### Cells and cell lines

Calu3 epithelial lung cancer cells were kindly provided by Laureano de le Vega, Dundee University, Scotland, UK. Cells were grown in Dulbecco’s Modified Eagle Medium (DMEM) (ThermoFisher) supplemented with 10% heat-inactivated FCS (Sigma-Aldrich), 1000 U ml−1 penicillin (Gibco), 1000 μg ml^−1^ streptomycin (Gibco), and 2 mM L-glutamine (Gibco). Cells were maintained at 37° with 5% CO_2_. A549 lung adenocarcinoma cells were originally obtained from German Collection of Microorganisms and Cell Cultures GmbH (DSMZ, Braunschweig, Germany) and were grown in DMEM medium supplemented with 10% FCS and 2 mM L-glutamine. Huh-7.5 and HEK293T cells were cultured in Dulbecco’s modified Eagle’s medium supplemented with 10% fetal calf serum, 2 mM L-glutamine, 0.1 mM nonessential amino acids, and 1% penicillin/streptomycin at 37°C and 5% CO_2_. Generation and verification of *NRF2^−/-^* iPSC was described in [43], and the absence of NRF2 gene expression was verified again by RT-qPCR (**Figure S5G**). Isogenic WT and *NRF2^−/-^*iPSC cells [43] were differentiated into CD31+ vascular ECs using an established protocol [44] and cultured in ECGM-2 medium (PromoCell, Darmstadt, Germany) on plates coated with fibronectin (Corning, NY, USA).

## Methods Aarhus (Figure 1C - G)

### Electroporation of Calu3 cells with gRNA and Cas9 protein

Calu3 CRISPR/Cas9 knockout cells were generated based on the recommendations of the manufacturer (Synthego) using the 4D-Nucleofector (Lonza) program CM-138. Briefly, cells were detached using TrypLe Express (Gibco) and cells were nucleofected with a complex of 24 μg of Cas9 Nuclease V3 protein (ID technology) + 12.8 μg of sgRNA (Synthego, USA) in OptiMEM media (Gibco). After nucleofection, cells were resuspended in prewarmed RPMI medium and incubated for 48 h until further experiments. Sequences of the synthetic guide RNA used are as follows; *AAVS1* (served as control) - G*G*G*GCCACUAGGGACAGGAU, *NFE2L2* - A*U*U*UGAUUGACAUACUUUGG.

### Short-interfering RNA (siRNA)-mediated knock down in Calu 3 cells

Calu3 cells were transfected in 6-well plates with 80 pmol of human Nrf2 (sc-37030) or control si RNA (sc-37007) (Santa Cruz Bio) diluted in serum and antibiotic free DMEM and using Lipofectamine RNAi Max as per manufacturer’s instructions. Calu 3 cells were incubated for 48 h in the presence of the siRNA before being processed.

### SARS-CoV-2 and variants of concern

SARS-CoV-2 Wuhan-like early European B.1 lineage (FR-4286) was kindly provided by Professor Georg Kochs (University of Freiburg). Professor Arvind Patel (University of Glasgow, UK) kindly provided the alpha variant B.1.1.7, and the beta variant B.1.351 was kindly provided by Professor Alex Sigal, African Health Research Institute, South Africa. Delta variant B.1.617.2 (SARS-CoV2/hu/DK/SSI-H11) was provided by Statens Serum Institut, SSI, Denmark. The viruses were propagated as previously described [45]. Briefly, 10 × 10^6^ VerohTMPRSS2 cells were seeded in T175 cell culture flasks and infected the following day at a MOI of 0.005 in 10 mL of cell culture media containing serum. After 1 h the cell culture medium was increased to 20 mL per flask and virus propagation continued for 72 h. Cell debris was removed by centrifugation of the cell culture supernatants from the different flasks at 300 g for 5 min. Viruses were concentrated in Amicon filter tubes by spinning at 4000 g for 30 min at 4°C. The concentrated virus was further aliquoted and stored at -80°C. The amount of infectious virus was determined using an endpoint dilution assay (TCID_50_) as described below.

### SARS-CoV-2 infection experiments

Calu3 cells were seeded in a 24-well plate and were allowed to adhere overnight. The respective compounds were added at the indicated concentrations either 24 or 48 h prior to viral infection. Cells were infected with SARS-CoV-2 in the presence of the compounds. After 1 h the medium was replaced with fresh DMEM containing the compounds.

### RT-qPCR for SARS-CoV-2

Cells were washed with PBS and lysed in 300 µL of RNA lysis buffer (Roche) diluted with 200 µL of PBS, followed by RNA extraction and gene expression analysis. RNA was extracted using the High Pure RNA Isolation Kit (Roche) according to the manufacturer’s instructions. RNA samples were diluted to 100 ng/uL and gene expression was analyzed by real-time quantitative PCR using TaqMan® RNA-to-CT ™ 1-Step Kit (Applied Biosystems) as previously described [45]. For SARS-CoV-2 gene detection, primers and probe sequences, hybridizing to the nucleocapsid region, were provided by the CDC and purchased from Eurofins. Samples were analyzed in a final volume of 10 μL, containing 5 μL of master mix, 0.5 μL (10 pmol/μL) of forward primer, 0.7 μL of reverse primer (10 pmol/μL), 0.2 μL of probe (20 pmol/μL), 2.4 μL of nuclease-free water, and 1 μL of diluted RNA. The analysis was performed on a ThermoFisher Scientific qPCR machine (Quant Studio 5). mRNA encoding TATA-Box Binding Protein (TBP) was used as internal control and was obtained from ThermoFisher.

### LDH release assay

Cellular toxicity of the respective compounds was measured 48h post-treatment using CyQUANT™ LDH Cytotoxicity Assay Kit (ThermoFisher) according to the manufacturer’s instructions. Untreated cells were used as a negative control, whereas cells lysed with the provided lysis buffer served as a positive control. LDH activity was determined following subtraction of the background 690 nm absorbance value from the 450 nm absorbance value measured on a BioTek Microplate Reader (BioTek Instruments). The percentage of cytotoxicity was calculated using the following formula: (LDH activity – LDH activity control) / (LDH activity positive control – LDH activity negative control) × 100.

### Immunoblotting

Immunoblotting was performed as described [46]. Briefly, cells were washed with PBS and subsequently lysed in 100 µL ice-cold Pierce RIPA lysis buffer (Thermofisher Scientific), supplemented with 10 mM NaF, 1x complete protease inhibitor cocktail (Roche) and 5 IU/mL Benzonase Nuclease (Sigma). Protein concentration was measured using a BCA Protein Assay Kit (Thermofisher Scientific). Whole-cell lysates were resuspended in a loading buffer, consisting of XT Sample Buffer (BioRad) and XT Reducing Agent (BioRad). Samples were then denatured at 95°C for 5 minutes (min). 10-40 μg of the reduced sample was separated by SDS-PAGE on a 4-20% Criterion TGX pre-cast gradient gels (BioRad). Each gel was first run at 70 V for 20 min, following 110 V for 50 min. Proteins were transfer onto a Midi Format 0,2 μM PVDF membrane (BioRad) using a Transblot Turbo Transfer System (BioRad) for 7 min. Membranes were blocked for 1 h at room temperature with 5% skim milk (Sigma-Aldrich) in PBS supplemented with 0.05% Tween-20 (PBS-T). Membranes were fractionated into smaller pieces and incubated overnight at 4°C with one of the following primary antibodies in PBS-T and 0.02% sodium azide: anti-SQSTM1/p62 (#8025, Cell Signaling 1:1000), anti-LC3B (#3868, Cell Signaling 1:1000), anti-SARS-CoV-2 Nucleocapsid (#26369, Cell Signaling, 1:1000) and anti-Vinculin (#V9264, Sigma 1:10.000) was used as a loading control. Membranes were washed 3 times in PBST for 15 min following incubation with secondary antibodies: HRP conjugated F(ab)2 donkey antirabbit IgG (H+L) or HRP conjugated F(ab)2 donkey anti-mouse IgG (H+L) (1:10.000) (Jackson ImmunoResearch) in PBS-T 1% skimmed-milk for 1h at RT. Membranes were washed 3 times in PBS-T for 10 min and subsequently incubated with SuperSignal West Dura Substrate or SuperSignal West Femto Maximum Sensitivity Substrate (ThermoFisher Scientific) for 1 min prior to exposure using iBright CL1500 Imaging System (ThermoFisher Scientific).

## Methods (Hannover)

### Viruses

ARS-CoV-2 infection assays were carried out using strain SARS-CoV-2/München-1.2/2020/984,p3) (Wölfel et al., 2020), kindly provided by Christian Drosten (Charité, Berlin) through the European Virus Archive – Global (EVAg). After primary isolation from the patient, the virus was propagated in and titrated on Vero cells (passage 3). The recombinant HCoV-229E encoding a *Renilla* luciferase gene (kindly provided by Volker Thiel, University of Bern, Switzerland, [47], here referred to as 229E-luc) was grown at 33°C on Huh7.5 cells and titrated in the same cell line.

### Infection assays

For SARS-CoV-2 infection, 4.5×10^5^ Calu-3 cells/well were seeded in 24-well plates coated with collagen. The next day, cells were pretreated with the indicated compounds for 24 h at 37°C and 5% CO_2_ and then inoculated with the SARS-CoV-2 isolate (MOI 0.005) in presence of the compounds and controls. Heat-inactivated virus (15 min at 70°C) served as input viral RNA (vRNA) control. The inoculum was removed after 4 h and cells washed twice with 1x PBS and fresh medium complemented with compounds. At 48 h p.i., vRNA was isolated from supernatants (NucleoSpin RNA kit, Macherey Nagel) and cell lysates (QIAamp Viral RNA Mini Kit, QiaGen) and viral genome copies were determined by RT-qPCR using SuperScriptIII one-step RT-PCR and Platinum Taq Polymerase (Invitrogen) [48]. vRNA amplification methodology was described previously [49]. For HCoV-229E-luc infections of iPSC-derived ECs and A549 cells, cells were seeded in 12-or 96-well plates and inoculated with a luciferase-labeled HCoV-229E-luc at MOI of 0.3 for 4 h. After removing the viral inoculum, fresh cell culture media complemented with compounds, was added and the cells were collected 48 h.p.i either for RT-qPCR to determine viral genome copies using SuperScript™ III One-Step RT-PCR System Platinum™ Taq DNA Polymerase (Thermo Fisher Scientific) or for *Renilla* luciferase activity to determine infectivity and replication.

### VSV pseudoparticle production and cell entry assay

Recombinant VSV pseudoparticles bearing SARS-CoV-2, SARS-CoV-1 or VSV glycoproteins were generated as described [50, 51]. In short, HEK293T cells were transfected with vectors expressing SARS-CoV-2-S, SARS-CoV-1-S, VSV-G, or with empty expression vector as negative control. After 24 h, cells were inoculated for 1 h with a replication-deficient VSV vector containing a Firefly luciferase and an eGFP expression cassette instead of the VSV glycoprotein (kindly provided by Gert Zimmer, Institute of Virology and Immunology, Switzerland). Subsequently, inoculum was removed, cells washed twice with 1x PBS, and medium supplemented with anti-VSV-G antibodies (I1, mouse hybridoma supernatant from CRL-2700) added, except to those viruses expressing VSV-G. Cell culture supernatant was harvested 16 h post inoculation and clarified from cell debris by centrifugation (2000 x g for 10 min) prior storage at -80°C. For cell entry assays, 5×10^4^ Calu-3 cells/well were seeded on collagen-coated 96-well plates. The next day, cells were pretreated with the indicated compound for either 24 h or 48 h, refreshing compounds after 24 h for the latter. After pretreatment, cells were washed with 1x PBS and inoculated with 50 µl of the indicated pseudoparticles in the absence of the compounds. 90 min post inoculation, 150 µl of complete media was added onto the inoculum. At 16 h post inoculation, cells were washed once with 1x PBS and lysed with 50 µL/well of Luc-lysis buffer (1% Triton-X 100, 25 mM Gly-Gly (pH 7.8), 15 mM MgSO4, 4 mM EGTA, 1 mM DTT in H_2_O). Transduction efficiency was measured by luciferase activity using a microplate reader (Berthold Technologies). To show SARS-CoV-1 and SARS-CoV-2 spike-specific entry effects, the background for each treatment was subtracted (values from the negative control; no env pseudoparticles), and data were normalized to the signal obtained with VSV-G.

### Gene knock-down with siRNA

For XPO1 knock-down, cells were grown to 90% confluency and transfected with specific (ON-TARGETplus Human XPO1 (7514) siRNA—SMARTpool, 5 nmol, L-003030-00-0005, Horizon Discovery) or control siRNA (ON-TARGETplus Non-targeting Control Pool, D-001810-10-05, Horizon Discovery) using Opti-MEM medium (31985070, Gibco). ON-TARGETplus Human ABCB1 siRNA—SMARTpool, 5 nmol (L-003868-00-0005, Horizon Discovery) was used to knock down *ABCB1* mRNA. ON-TARGETplus Human HMOX1 siRNA—SMARTpool, 5 nmol, (L-006372-00-0005, Horizon Discovery) was used to knock down*HMOX1* mRNA. Knock-down of XPO1 mRNA and protein was verified after 24 h by RT-qPCR and immunoblot, respectively. Knock-down of *ABCB1* and *HMOX1* mRNA was verified after 24 h by RT-qPCR.

**Real-time quantitative reverse transcriptase polymerase chain reaction (RT-qPCR)** of host cell mRNA was performed using Nucleospin RNA purification kit (Machery Nagel), on-column removal of DNA with rDNase (Machery Nagel), and the PrimeScript cDNA synthesis kit (TaKaRa, Shiga, Japan) with 400 ng RNA input in a 10 µL reaction. Sequences of PCR primers are shown in **Table S2**. A LightCycler® 2.0 instrument (Roche, Mannheim, Germany) was used, employing 45 thermocycles of 95°C for 15 sec., 60°C for 15 sec., and 72°C for 15 sec. Melting curve analysis was performed to exclude artifacts resulting from primer dimer formation, using the sequence 95°C for 15 sec., 60°C for 15 sec., 95°C for 1 min. and 37°C for 30 sec. Relative expression of target mRNAs was calculated using the 2^−ΔΔCT^ method [45], using *HPRT* mRNA as internal reference.

### Assessment of *XPO1* mRNA dynamics in the presence of ActD and 4OI

A549 cells were seeded in 12-well plates to a density of 2×10^5^ cells/well. ActD (A9415, Sigma Aldrich) and/or 4OI were added to the medium to concentrations of 5 µg/ml and 100 µM, respectively. As indicated in the figure legend, ActD-containing medium was replaced after 24 h with fresh medium containing neither compound or 100 µM 4OI. XPO1 mRNA levels were measured by RT-qPCR after 48 h.

**Immunoblotting** was performed as described in detail in [52], using a semi-dry transfer system (Trans-Blot Turbo, BioRad), Amersham enhanced chemiluminescence western blot detection reagent (GE Healthcare Science, Pittsburgh, PA), and a Vilber fusion FX7 device (Vilber Smart Imaging, Collégien, France). The following primary antibodies were used: ACE2 (92485S, 1:1000, Cell Signaling Technology), XPO1 (46249S, 1:1000, Cell signaling technology), β-actin (ab49900, 1:20,000, Abcam). Goat anti-rabbit IgG-HRP (Southern Biotech, catalogue no. 4030– 05) was used as secondary antibody.

### Mitochondrial ROS assay

229E-luc infection (MOI = 0.3) and treatments were carried out as described above. Upon conclusion of the experiment, the cells were incubated for 5 min with medium containing 5 μM MitoSOX Red mitochondrial superoxide indicator (Invitrogen, cat# M36008). After washing with PBS, cells were resuspended with cold PBS for measurement of mitochondrial ROS by flow cytometry (Sony SP6800 ZE Analyzer, phycoerythrin channel).

### RNA sequencing

Quality and integrity of total RNA was controlled on Agilent Technologies 2100 Bioanalyzer (Agilent Technologies; Waldbronn, Germany), using only samples with RNA Integrity Number (RIN) of ≥8 for subsequent RNA sequencing. Libraries were prepared from 500 ng total RNA using Dynabeads® mRNA DIRECT™ Micro Purification Kit (Thermo Fisher) for mRNA purification, followed by the NEBNext® Ultra™ II Directional RNA Library Prep Kit (New England BioLabs) according to the manufactureŕs protocols. The libraries were sequenced on an Illumina NovaSeq 6000 using the NovaSeq 6000 S1 Reagent Kit (100 cycles, paired-end run) yielding an average of 5 ×10^7^ reads per RNA sample. Quality reports for each FASTQ file were generated using the FASTQC tool. Before alignment to the reference genome, raw sequences were trimmed based on base call quality and adapter contamination using fastq-mcf. Reads shorter than 15 bp were removed from the FASTQ file. Trimmed reads were aligned to the reference genome (hg38) using the open source aligner STAR (https://code.google.com/p/rna-star/) [53] with settings recorded in the log file. Raw counts were generated with the R package **RSubread** [54], and features were annotated using **biomaRt** (https://www.ensembl.org/info/data/biomart/index.html). Features categorized as “rRNA” or “pseudogene” were removed from the dataset. Only features with more than 3 counts per million (CPM) in at least one set of replicates were included. Data normalization was performed using the TMM (Trimmed Mean of M-values) method [55], and normalized expression values (CPM) were log2-transformed before differential expression analysis.

### Bioinformatics

RNAseq data were analyzed using DESeq2 (differential expression analysis, Venn diagrams, hierarchical clustering analysis and KEGG pathway GSEA). Metaboanalyst (https://www.metaboanalyst.ca/) [56] was used to prepare heatmaps. The strip charts (**Figure 7**, **Figure S7**) were generated using R package ggplot2. The GSEA of XPO1 cargoes was performed with data from [20], using Webgestalt (https://www.webgestalt.org/). Network analysis of predicted E3 ligase targets was performed using UbiBrowser (http://ubibrowser.bio-it.cn/ubibrowser/home/index).

### Statistics

Data were analyzed using GraphPad Prism v8.02 (GraphPad Software), which was also used to generated most graphs. Unless stated otherwise, statistical significance was determined by ANOVA with Tukey’s post-hoc test to correct for multiple hypothesis. Experiments were performed using three biological replicates unless stated otherwise in the figure legends. Data are shown as means +/-standard deviation (SD).

## Acknowledgments

We thank Lothar Jänsch (Helmholtz Centre for Infection Research, HZI) for helpful discussion and Michael Jarek and the staff of the HZI Genome Analytics group for expert RNA sequencing.

